# Solu – a cloud platform for real-time genomic pathogen surveillance

**DOI:** 10.1101/2024.05.30.596434

**Authors:** Timo J Moilanen, Kerkko Visuri, Jonatan Lehtinen, Irene Ortega-Sanz, Jacob L Steenwyk, Samuel Sihvonen

## Abstract

**Background:** Genomic surveillance is extensively used for tracking public health outbreaks and healthcare-associated pathogens. Despite advancements in bioinformatics pipelines, there are still significant challenges in terms of infrastructure, expertise, and security when it comes to continuous surveillance. The existing pipelines often require the user to set up and manage their own infrastructure and are not designed for continuous surveillance that demands integration of new and regularly generated sequencing data with previous analyses. Additionally, academic projects often do not meet the privacy requirements of healthcare providers.

**Results:** We present Solu, a cloud-based platform that integrates genomic data into a real-time, privacy-focused surveillance system.

**Evaluation:** Solu’s accuracy for taxonomy assignment, antimicrobial resistance genes, and phylogenetics, was comparable to established pathogen surveillance pipelines. In some cases, Solu identified antimicrobial resistance genes that were previously undetected. Together, these findings demonstrate the efficacy of our platform.

**Conclusions:** By enabling reliable, user-friendly, and privacy-focused genomic surveillance, Solu has the potential to bridge the gap between cutting-edge research and practical, widespread application in healthcare settings. The platform is available for free academic use at platform.solugenomics.com.

## Background

Bacterial and fungal pathogens, along with their antimicrobial resistance, are causing an increasing burden on healthcare and public health (1–3). Advances in microbial genomics have significantly enhanced infection prevention and outbreak surveillance by providing detailed information about pathogen species, antimicrobial resistance, and phylogenetics (4,5). As the cost of Whole-Genome Sequencing (WGS) has decreased rapidly, continuous genomic surveillance has become a cost-effective method for infection prevention and control (6,7). The interest towards genomic analysis has led to the emergence of several pathogen analysis tools, such as nf-core (8), TheiaProk (9), ASA3P (10), CamPype (11), Nullarbor (12), Bactopia (13), and Galaxy (14), which enable genomic analysis also for users without in-depth expertise in bioinformatics or computer science.

Despite these advancements, bioinformatics still remains a bottleneck for the widespread adoption on pathogen genomic surveillance due to limitations in usability, speed, and security (7).

Most existing pipelines (8–13) are operated using the command-line interface (CLI) and require the user to manage their own data storage and computation infrastructure. While it is possible to learn their usage without advanced computational knowledge (15), many practitioners simply don’t have the time or willingness for it and prefer graphical user interfaces instead. Additionally, most existing tools are designed for single-use execution, which is a challenge for continuous surveillance where new sequencing data is often generated in small batches (6,16). To facilitate ongoing analysis, users must implement their own processes for integrating new and old data.

Fast time to results has been identified as a key component for effective genomic surveillance (6,16). As new samples arrive in batches and need to be compared to all previously accumulated samples, computation time can become a significant bottleneck if using a single-workstation installation. Also, unless the pipelines are highly automated, running the analyses often requires specially trained personnel who might not be available immediately upon the arrival of new data.

Academia-led projects developed under FAIR (Findable, Accessible, Interoperable, Reusable) principles often lack the necessary privacy focus to meet the stringent requirements of healthcare providers (17). In contrast, healthcare providers must adhere to stringent legal requirements, such as the U.S. HIPAA Privacy Rule (18), ruling out many existing online platforms for genomic surveillance.

It is possible for healthcare providers to overcome these limitations by implementing their own automated pipelines, but it requires significant investments in bioinformatics and computational infrastructure, and the lack of these resources is a challenge in many facilities (19). To fill this gap, we present Solu – an automated, fast, and secure web application for analyzing WGS samples.

## Implementation

Solu is a cloud-based platform for the analysis of bacterial and fungal WGS samples. Its automated bioinformatics pipeline includes genomic characterization and phylogenetic comparison. Its cloud implementation is built to match the usability, speed, and security requirements of ongoing genomic surveillance in healthcare facilities.

### Bioinformatics pipeline

The platform runs a fully automated pathogen analysis pipeline, which is illustrated in Figure 1. The pipeline includes *de novo* assembly, quality assurance (QA), species identification and genomic characterization for each uploaded sample, and phylogenetic comparison between all uploaded samples of the same species. It is triggered automatically after each file upload and cannot be configured by the user. This section presents an overview of the pipeline, and a detailed description can be found in Supplementary Material 1.

**Figure 1.**
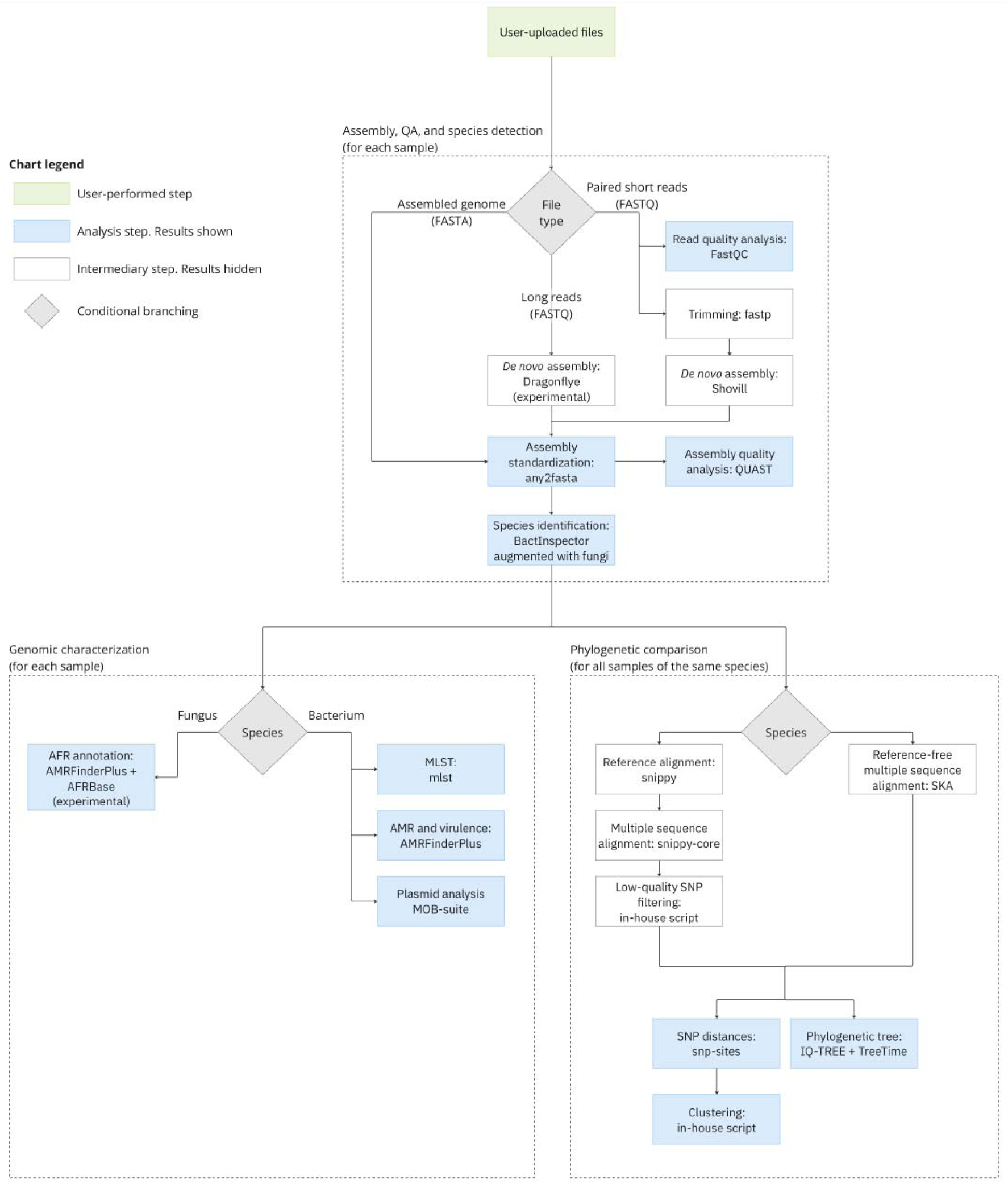
Bioinformatics pipeline.

### Input format

The supports three input types: paired-end short reads in FASTQ format, long reads in FASTQ format, or an assembled genome in FASTA format. Analysis of long reads is still considered an experimental feature.

### Assembly, QA, and species detection

Short reads are quality checked using FastQC (20), quality corrected with fastp (21), and assembled using Shovill (22). Long reads are pre-processed, assembled and polished with Dragonflye (23). After assembly, all samples are standardized using any2fasta (24) and quality assessed with Quast (25).

Species is identified with Bactinspector (26). To identify fungal species, Bactinspector’s default database was augmented with all fungal reference genomes from the NCBI Taxonomy (27). The augmented database also includes clade-level reference genomes for *Candida auris*.

### Genomic characterization

Analysis of bacterial species includes multi-locus sequence typing (MLST) with mlst (28), AMR annotation using AMRFinderPlus (29), and plasmid analysis with MOB-suite (30).

The pipeline also includes an experimental antifungal resistance (AFR) gene annotation for the species *Candida auris*. AFR annotation is implemented using AMRFinderPlus with a custom database of known AFR point mutations sourced from AFRBase (31).

### Phylogenetic comparison

The pipeline’s phylogeny is based on constructing a multiple sequence alignment for each species. Based on the species in question, multiple sequence alignment is computed by either a reference-based or reference-free method.

The reference-based alignment is considered more robust and has been implemented for 21 commonly analyzed species. It includes aligning each sample to the species’ reference genome using Snippy (32), creating a multiple-sequence alignment using snippy-core (32), and filtering out low quality SNPs using an in-house script.

The reference-free alternative is implemented to support analysis of species that are not yet supported by the reference-based alignment. It is computed using the split-kmer analysis tool SKA (33).

After constructing the multiple sequencing alignment, the phylogenetic comparison includes pairwise SNP distances, clustering, and phylogenetic tree inference. SNP distances are counted from the multiple sequence alignment using snp-sites (34) and snp-dists (35).

Samples are clustered with a 20-SNP single-linkage clustering threshold using an in-house Python script. Phylogenetic trees are inferred using a general time reversible maximum likelihood model from IQ-TREE 2 (36) and midpoint-rooted using TreeTime (37). Both IQ-TREE 2 and TreeTime are run using the Augur toolkit (38).

### Automated cloud infrastructure

The cloud infrastructure of Solu is built on three principles: usability, speed, and security.

### Usability

The Solu Platform is web based, enabling practitioners to use it without installing software or running command-line tools (Figure 2). New samples are uploaded using the drag-and-drop web UI, which automatically triggers a bioinformatics pipeline. The pipeline requires zero configuration from the user, which promotes repeatability and alleviates the need for in-depth bioinformatics knowledge. The analysis results are stored in the cloud, eliminating the need for a self-implemented storage system and enabling effortless result sharing between colleagues.

**Figure 2.**
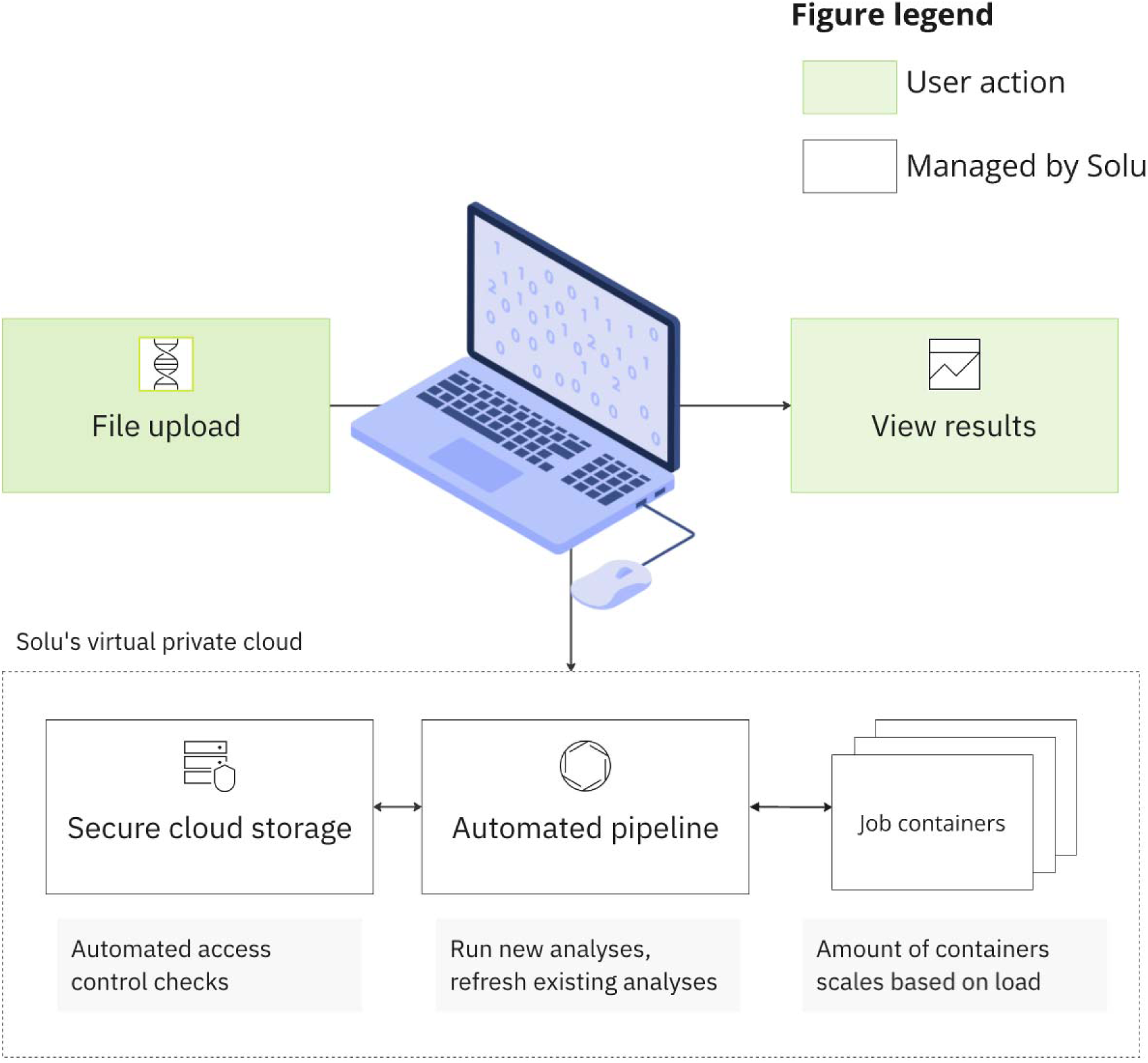
Solu’s cloud platform implementation.

Each newly uploaded sample is also automatically compared to previously uploaded samples of the same species, which enables detecting potential new outbreaks quickly. For instance, when a user uploads a *Salmonella enterica* sample, the platform automatically re-computes the phylogenetics for all uploaded *Salmonella enterica* samples and highlight possible clusters.

### Speed

The platform’s cloud infrastructure is optimized for speed even during peak usage. This is achieved by running each computation-intensive workload in a separate Docker container with optimized resource (CPU and memory) distribution. These containers are orchestrated by a cloud computing cluster that is automatically scaled up and down based on usage, up to a maximum of 512 CPUs and 2 TB of memory. The cluster also contains a pool of hot standby resources, which allows starting the analysis of a new sample within seconds of its upload.

This auto-scaling capability brings the user substantial speed improvements by allowing the parallelization of some of the analyses in the pipeline. In addition, it allows analyzing a whole batch of samples simultaneously, leading to a significant reduction in overall time-to-results when analyzing a batch of samples. Importantly, these speed improvements are achieved without a significant increase in computation costs.

### Security

The platform’s data is stored in a secure cloud storage with set read and write permissions. All computations occur within a virtual private network, monitored by automated access control checks. Solu implements strict data security protocols, including appropriate access permissions, encryption, continuous monitoring, code reviews, staff training, and other cybersecurity measures. Accordingly, Solu adheres to the U.S. HIPAA rule, and can sign a Business Associate Agreement (BAA) for enterprise customers. Solu also allows enterprise customers to choose between U.S. or EU as their data storage. Further information regarding data security practices can be found at https://solugenomics.trust.site/.

### Evaluation

To evaluate the Solu platform, we reproduced four outbreak investigation studies using published genomic data (**Table 1**). Data was obtained from the European Nucleotide Archive as raw reads and uploaded to the Solu platform. All samples were paired-end short-reads in FASTQ format.

**Table 1.** Overview of evaluation datasets.

| Species | BioProject accession | Number of samples | Raw data size (GB, zipped) |
| --- | --- | --- | --- |
| *Staphylococcus aureus* | PRJNA400143 (40) | 135 | 16.47 |
| *Enterococcus faecium* | PRJEB34664 (41) | 99 | 15.18 |
| *Salmonella enterica* | Multiple (42) | 23 | 5.03 |
| *Candida auris* | PRJNA328792 (43) | 47 | 50.94 |

We evaluated Solu’s performance by computing metric scores for species identification, MLST, clade construction, and AMR predictions against the references. Phylogenetic trees were exported from Solu. Tree topologies were compared visually, and where raw tree data was available, we calculated a Robinson-Foulds distance using TreeDist (39). Both reference-based and reference-free phylogenetic pipelines were run for all datasets and compared against each other to validate the platform’s internal consistency.

We also measured the required time for analysis of each sample. Plasmids were not evaluated in this study due to the absence of plasmid annotations in the original studies.

## Results

### Evaluation results

The Solu platform successfully completed the bioinformatics pipeline for all 304 samples. A screenshot of the platform’s home screen is shown in Figure 3. This workspace, including all samples and results, is also accessible at a user-friendly web interface at https://platform.solugenomics.com/w/solu-publication-2 and https://platform.solugenomics.com/w/solu-publication-3.

**Figure 3.**
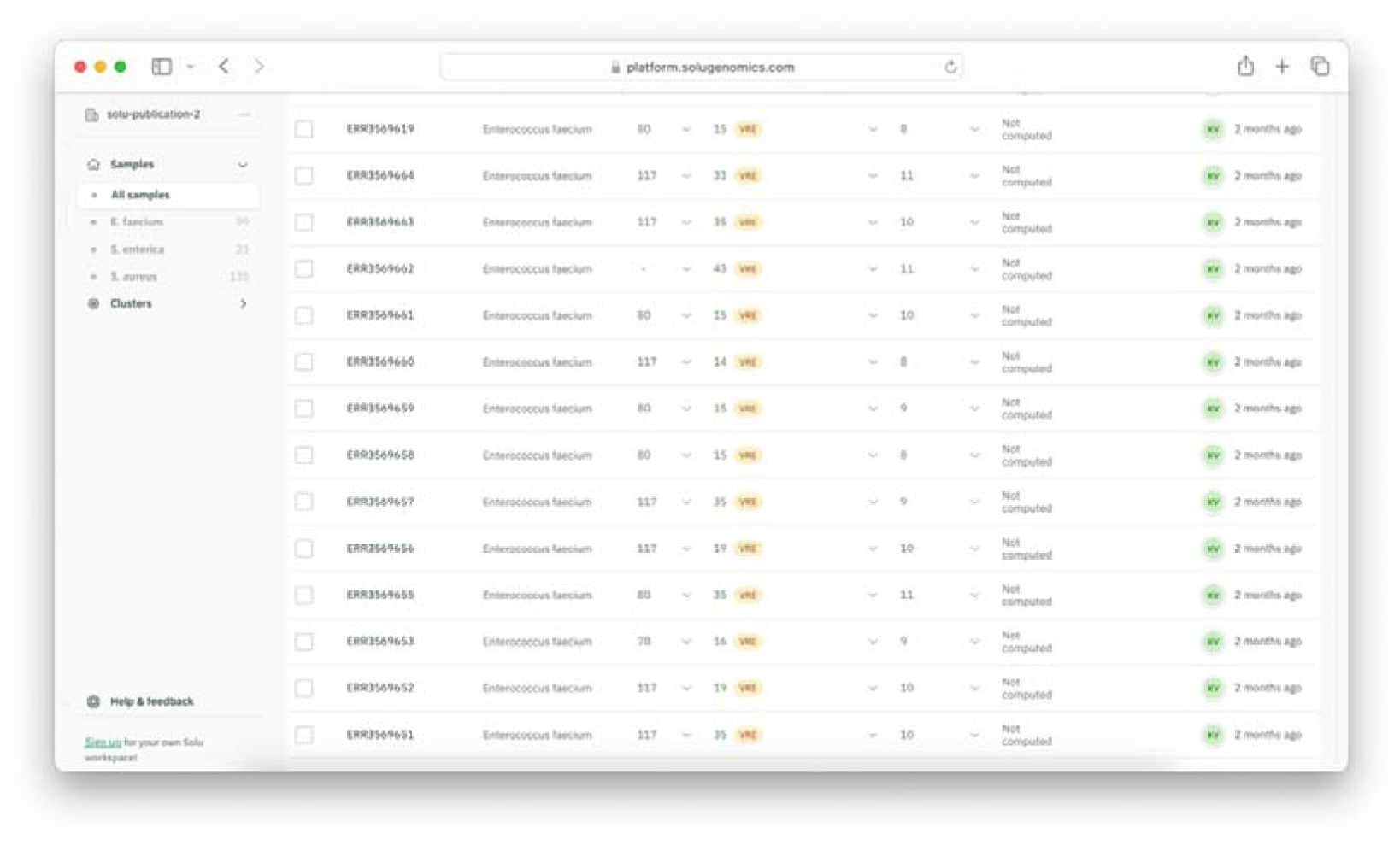
Summary of the samples shown in Solu.

### Species, MLST and clade assignment

Solu accurately identified the species of all 304 samples and assigned the correct clade for all 47 *Candida auris* samples.

Exact MLST matches were observed in 210 out of 230 (91.3%) isolates with known sequence types (Table 2). However, 18 of 20 non-exact MSLT matches were single-locus variants.

**Table 2.**
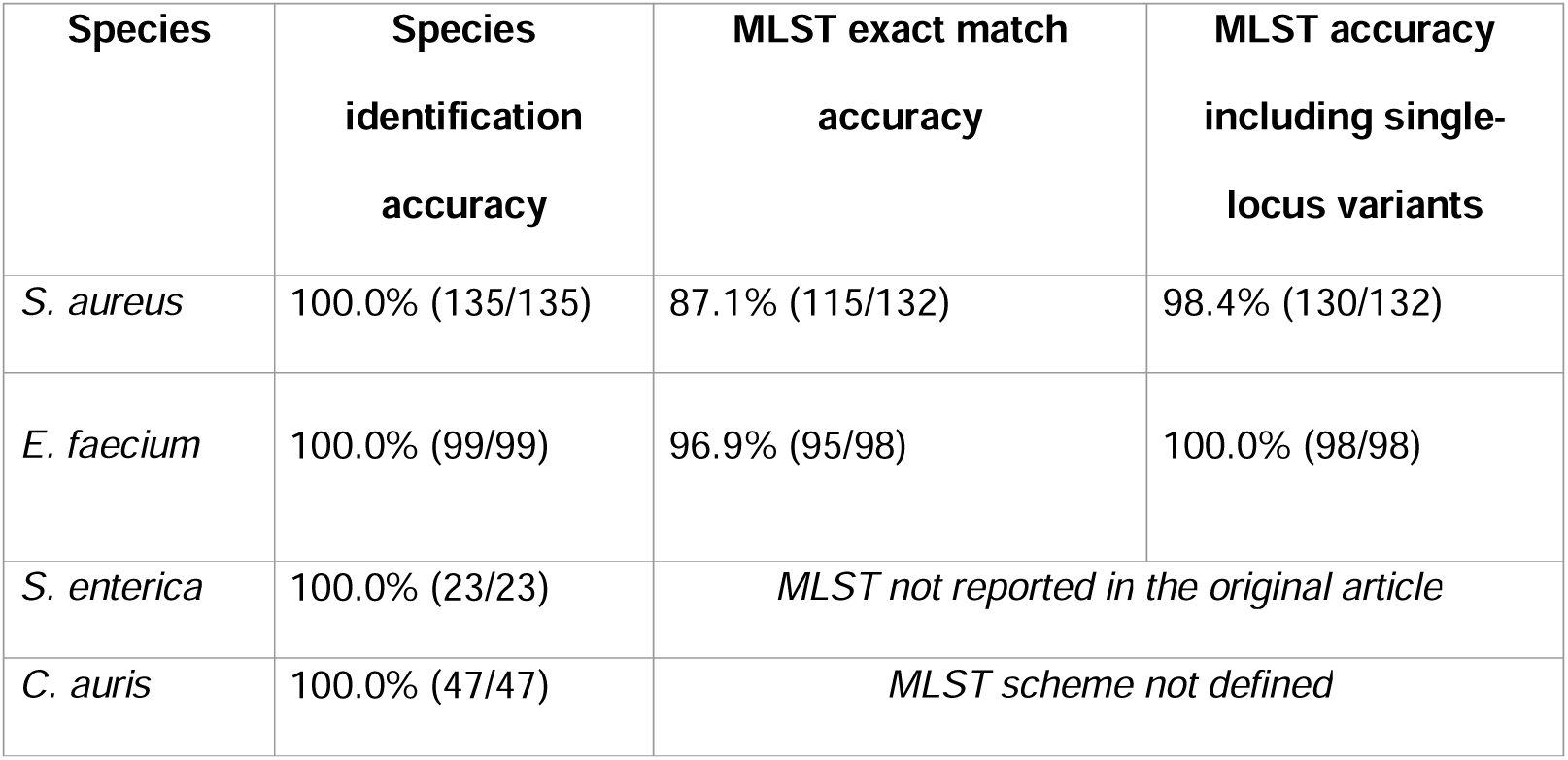
Species identification and MLST concordance. Solu’s results compared to the original publications when available there.

| Species | Species identification accuracy | MLST exact match accuracy | MLST accuracy including single-locus variants |
| --- | --- | --- | --- |
| *S. aureus* | 100.0% (135/135) | 87.1% (115/132) | 98.4% (130/132) |
| *E. faecium* | 100.0% (99/99) | 96.9% (95/98) | 100.0% (98/98) |
| *S. enterica* | 100.0% (23/23) | *MLST not reported in the original article* |  |
| *C. auris* | 100.0% (47/47) | *MLST scheme not defined* |  |

### Antimicrobial resistance

The antimicrobial resistance (AMR) gene detection results from Solu were compared with those of the references. Concordance varied by species, ranging from 99.6% for *E. faecium* to 93.1% for *S. aureus*. Table 3 summarizes some commonly studied AMR loci, while the full results can be viewed online.

**Table 3.** Sensitivity of Solu’s AMR prediction.

| Species | Overall AMR<br>locus<br>sensitivity | *vanA* | *vanB* | *mecA* | Other key genes |
| --- | --- | --- | --- | --- | --- |
| *S. aureus* | 93.1%<br>(309/332) | N/A | N/A | 98.8% (81/82) | *mupA*: 100%<br>(3/3),<br>*blaZ*: 90.9%<br>(100/110) |
| *E. faecium* | 99.6%<br>(515/517) | 100.0%<br>(30/30) | 100.0%<br>(69/69) | N/A | *ermB*: 100.0%<br>(95/95)<br>*tet*: 100.0% (4/4) |
| *S. enterica* | AMR results not reported in the original article |  |  |  |  |
| *C. auris* | 90.9% (40/44) | N/A | N/A | N/A | *ERG11_K143R*:<br>100.0% (8/8) |

Note: N/A indicates that the gene is not typically relevant for the species.

Key AMR genes, such as the *vanA/vanB* type and *mecA*, were detected with 100% and 98.8% sensitivity, respectively. Antifungal resistance mutation detection for Candida auris showed a 90.9% sensitivity. The results matched the original findings for 43 isolates.

However, Solu identified the *ERG11_K143R* mutation in 2 Clade I isolates, which were originally reported as having the *Y123F* mutation, and detected 2 isolates lacking ERG mutations.

The main reason for lower agreement in the *S. aureus* dataset was Solu’s inability to find any *dfrA* matches, whereas the original article reported 10 isolates with *dfrA*. (44)

### Phylogenetics

Solu automatically generated phylogenetic trees for all four datasets, which can be viewed and downloaded in the published workspace in “Tree view” (Figure 4).

**Figure 4.**
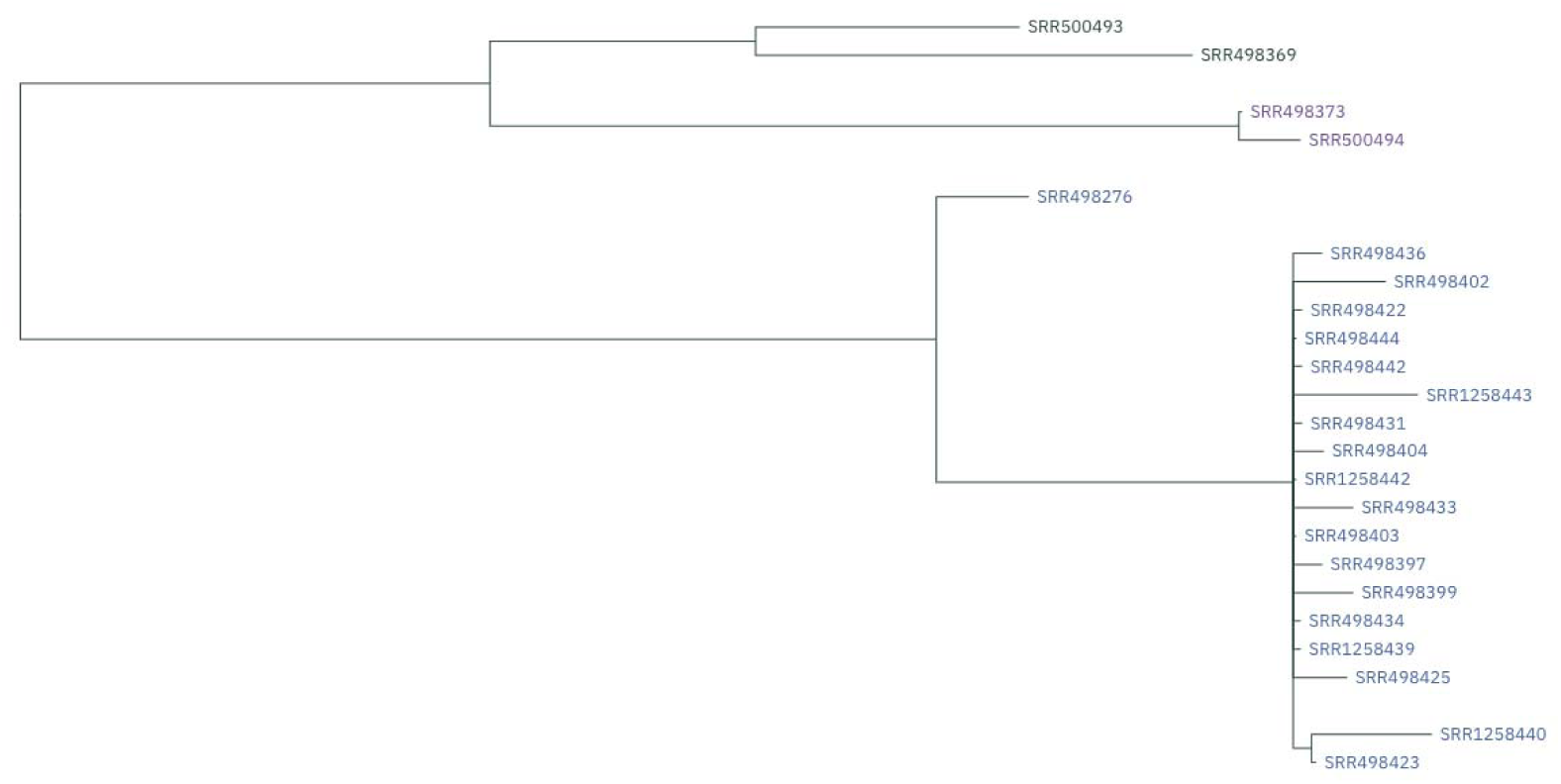
Screenshot of the *Salmonella enterica* tree in the graphical user interface.

The *E. faecium* phylogenetic tree generated by Solu demonstrated a high degree of concordance with the reported SNP subclusters, complex types (CTs) and sequence types (STs) of the reference (Figure 5). For the *Salmonella enterica* dataset, Solu produced a similar topology to the reference tree (Figure 6) where the outbreak samples are separate from the outgroup. Robinson-Foulds distance to the *S. enterica* reference tree was 2.

**Figure 5.**
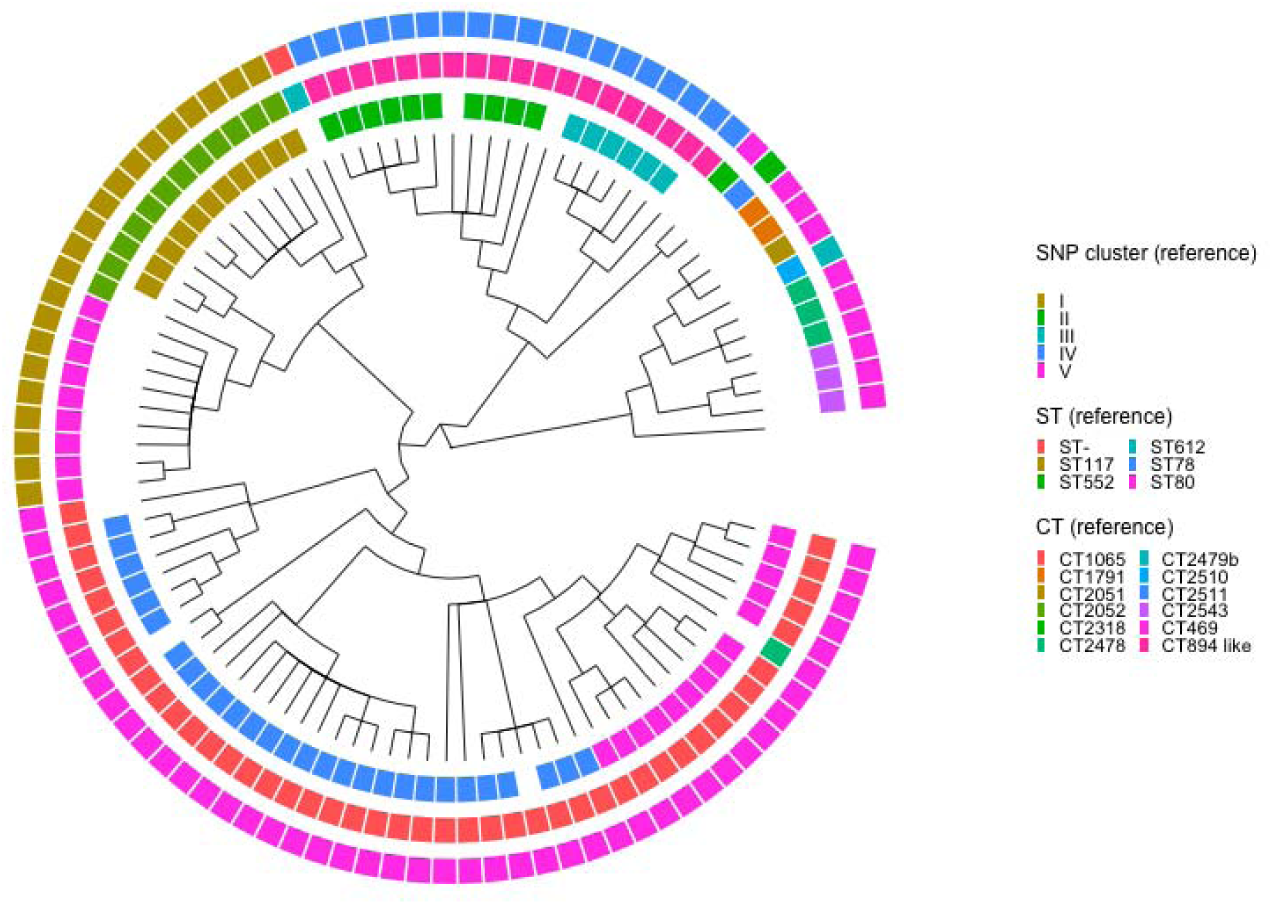
Solu’s *E. faecium* phylogenetic tree shown as a cladogram (inner) vs. SNP clusters, complex types (CT), and sequence types (ST) from the original publication (outer rings). The cladogram visualization was created using Dendroscope (45).

**Figure 6.**
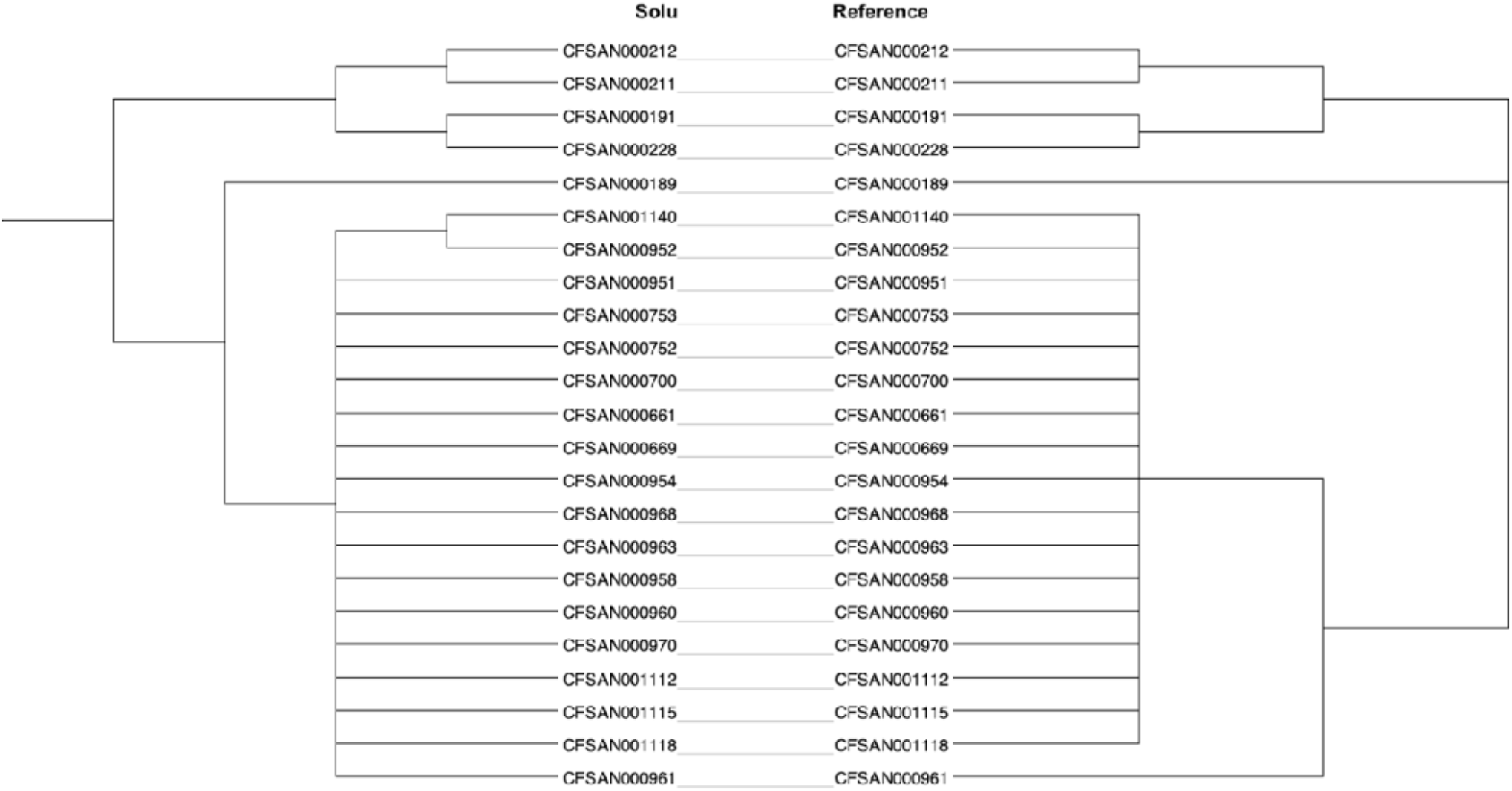
Tanglegram of Solu’s *Salmonella enterica* phylogenetic tree (left) vs. the reference tree (right).

For the *S. aureus* and *C. auris* datasets, Solu generated phylogenetic trees in which isolates with identical sequence types or clades consistently clustered together. Further detailed comparisons were not possible due to the lack of raw tree data and subtyping information.

The resulting trees are provided in Supplementary Material 2.

In comparing the reference-free and reference-based pipelines, Solu’s reference-free pipeline generated highly concordant phylogenetic trees with the reference-based pipeline. The Robinson-Foulds distances ranged from 0.08 to 0.46, as computed using TreeDist (46), indicating a high level of similarity (see Supplementary Material 2)

### Time-to-results

Time-to-results for the four datasets are presented in Table 4 and Figure 7. For bacterial samples, Solu completed de novo assembly in an average of 7.2 minutes and variant calling in 9.5 minutes from upload. For *C. auris* samples, de novo assembly averaged 17 minutes, and variant calling took 23.6 minutes from upload.

**Figure 7.**
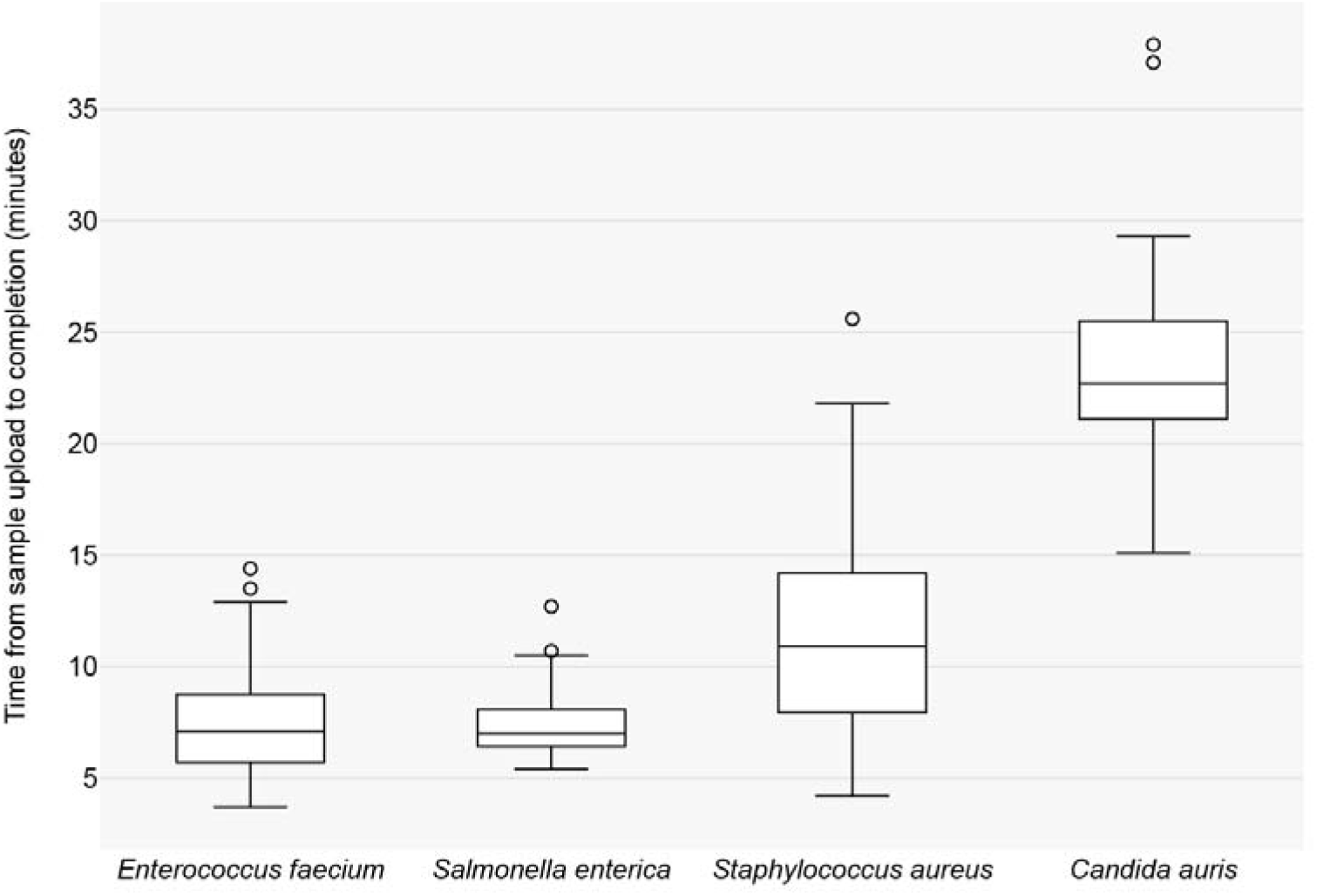
Box plot of Solu’s total time-to-results for each dataset.

**Table 4.** Average time-to-results per dataset. Shortest and longest recorded times in parentheses. Genome size. (**27**) **and sample count of each dataset included for additional context.**

|  | Genome size (Mb) | Sample count | Average total time from sample upload (minutes) |  |  |
| --- | --- | --- | --- | --- | --- |
|  |  |  | Read quality analysis and correction | De novo assembly | Variant calling |
| *Enterococcus faecium* | 2.9 | 99 | 1.5 (1.0–2.8) | 5.2 (1.9–11.8) | 7.4 (3.7–14.4) |
| *Salmonella enterica* | 5.0 | 23 | 1.4 (1.1–1.8) | 5.0 (2.9–10.1) | 7.6 (5.4–12.7) |
| *Staphylococcus aureus* | 2.8 | 135 | 1.9 (1.0–6.1) | 9.0 (2.4–21.9) | 11.4 (4.2–25.6) |
| *Candida auris* | 12.2 | 47 | 3.5 (2.4–5.9) | 17.0 (10.4–32.3) | 23.6 (15.1–37.9) |

## Discussion

Bioinformatics remains a bottleneck to widespread use of genomic pathogen surveillance. Usability, speed, and security are additional requirements for practical outbreak analysis in a healthcare setting.

Our evaluation demonstrates that the Solu platform produces outputs that are largely consistent with prior outbreak studies, using raw sequencing reads and requiring zero configuration, with a runtime of approximately 10 minutes for bacterial samples and 20 minutes for fungal samples. Solu’s phylogenetic pipelines produced results that were internally and externally consistent.

The largest discrepancies were observed in the *Staphylococcus aureus* dataset, where the original study used PCR for MLST assignment and applied 90% identity and 75% coverage thresholds for AMR gene detection. We hypothesize that the different pipeline parameters allow for higher sensitivity, at the cost of potential misidentification.

Compared to some other pipelines, Solu platform’s zero-configuration design prevents users from customizing pipeline parameters, which may result in some variation in the results. This approach was chosen to promote usability and prevent users from inadvertently selecting unsuitable parameters. Despite this limitation, default tool configurations provide sufficient accuracy for a wide variety of research applications, including AMR gene characterization and clonality assessment (47). Future studies leveraging more in-depth datasets and epidemiologically validated outbreaks hold great potential to further strengthen and expand the applicability of our findings.

We aim to improve the analytical capacity of the platform in future iterations, featuring additional tooling, modifications to the analytical workflow, broader support for species and databases, and improved runtimes among other features. We encourage users to contact the authors to request any additional analyses or databases of interest.

In conclusion, by focusing on a robust, privacy-focused infrastructure, Solu facilitates broader adoption of genomic pathogen surveillance, potentially bridging the gap between research and practice.

## Availability and requirements

Project name: Solu Platform

Project home page: https://platform.solugenomics.com

Operating system(s): Platform independent

Programming language: Typescript

Other requirements: Modern internet browser such as Google Chrome, Mozilla Firefox or Safari

Pricing, academic free plan, and non-academic license descriptions: www.solugenomics.com/pricing

## List of abbreviations

AFR: antifungal resistance; AMR: antimicrobial resistance; CPU: Central Processing Unit; CT: complex type; FAIR: Findable, Accessible, Interoperable, Reusable; HIPAA: Health Insurance Portability and Accountability Act; MLST: Multi-locus Sequence Typing; QA: quality assurance; SNP: single nucleotide polymorphism; ST: sequence type; WGS: Whole-Genome Sequencing.

## Declarations

### Ethics approval and consent to participate

Not applicable.

### Consent for publication

Not applicable.

### Availability of data and materials

The data analyzed in the study are available from the European Nucleotide Archive at https://www.ebi.ac.uk/ena using the accessions provided in Table 1. The Solu workspace, including all samples and results, is accessible at https://platform.solugenomics.com/w/solu-publication-2 and https://platform.solugenomics.com/w/solu-publication-3

## Authors’ contributions

TM: Conceptualization, Software, Formal Analysis, Investigation, Project Administration, Writing. KV: Software, Visualization, Methodology, Validation, Writing. JL: Software, Methodology, Validation, Writing. IOS: Methodology, Review & Editing, Visualization. JLS: Writing -- Review & Editing, Funding Acquisition, and Visualization. SS: Conceptualization, Writing. All authors read and approved the final manuscript.

## Acknowledgements

The authors thank the anonymous reviewers for their constructive comments.

## References

1. Murray CJL, Ikuta KS, Sharara F, Swetschinski L, Robles Aguilar G, Gray A, et al. Global burden of bacterial antimicrobial resistance in 2019: a systematic analysis. The Lancet. 2022 Feb;399(10325):629–55.

2. Vallabhaneni S, Mody RK, Walker T, Chiller T. The Global Burden of Fungal Diseases. Infect Dis Clin. 2016 Mar 1;30(1):1–11.

3. Van Rhijn N, Arikan-Akdagli S, Beardsley J, Bongomin F, Chakrabarti A, Chen SCA, et al. Beyond bacteria: the growing threat of antifungal resistance. The Lancet. 2024 Sep;404(10457):1017–8.

4. Eyre DW. Infection prevention and control insights from a decade of pathogen whole-genome sequencing. J Hosp Infect. 2022 Apr;122:180–6.

5. Steenwyk JL, Rokas A, Goldman GH. Know the enemy and know yourself: Addressing cryptic fungal pathogens of humans and beyond. Chowdhary A, editor. PLOS Pathog. 2023 Oct 19;19(10):e1011704.

6. Forde BM, Bergh, Haakon, Cuddihy, Thom, Hajkowich, Kristin, Hurst, Trish, Playford, E, et al. Clinical Implementation of Routine Whole-genome Sequencing for Hospital Infection Control of Multi-drug Resistant Pathogens. Clin Infect Dis. 2023 Feb 1;76(3):e1277–84.

7. Fox JM, Saunders NJ, Jerwood SH. Economic and health impact modelling of a whole genome sequencing-led intervention strategy for bacterial healthcare-associated infections for England and for the USA. Microb Genomics [Internet]. 2023 Aug 9 [cited 2024 Sep 17];9(8). Available from: https://www.microbiologyresearch.org/content/journal/mgen/10.1099/mgen.0.001087

8. Ewels PA, Peltzer A, Fillinger S, Patel H, Alneberg J, Wilm A, et al. The nf-core framework for community-curated bioinformatics pipelines. Nat Biotechnol. 2020 Mar;38(3):276–8.

9. Libuit KG, Doughty EL, Otieno JR, Ambrosio F, Kapsak CJ, Smith EA, et al. Accelerating bioinformatics implementation in public health. Microb Genomics. 2023 Jul 10;9(7).

10. Schwengers O, Hoek A, Fritzenwanker M, Falgenhauer L, Hain T, Chakraborty T, et al. ASA3P: An automatic and scalable pipeline for the assembly, annotation and higher-level analysis of closely related bacterial isolates. Pertea M, editor. PLOS Comput Biol. 2020 Mar 5;16(3):e1007134.

11. Ortega-Sanz I, Barbero-Aparicio JA, Canepa-Oneto A, Rovira J, Melero B. CamPype: an open-source workflow for automated bacterial whole-genome sequencing analysis focused on Campylobacter. BMC Bioinformatics. 2023 Jul 20;24(1).

12. Seemann T. nullarbor [Internet]. [cited 2024 Sep 17]. Available from: https://github.com/tseemann/nullarbor

13. Petit RA, Read TD. Bactopia: a Flexible Pipeline for Complete Analysis of Bacterial Genomes. Segata N, editor. mSystems. 2020 Aug 25;5(4).

14. Jalili V, Afgan E, Gu Q, Clements D, Blankenberg D, Goecks J, et al. The Galaxy platform for accessible, reproducible and collaborative biomedical analyses: 2020 update. Nucleic Acids Res. 2020 Jul 2;48(W1):W395–402.

15. Carriço JA, Rossi M, Moran-Gilad J, Van Domselaar G, Ramirez M. A primer on microbial bioinformatics for nonbioinformaticians. Clin Microbiol Infect. 2018 Apr;24(4):342–9.

16. Roberts LW, Forde BM, Hurst T, Ling W, Nimmo GR, Bergh H, et al. Genomic surveillance, characterization and intervention of a polymicrobial multidrug-resistant outbreak in critical care. Microb Genomics. 2021 Mar 1;7(3).

17. Stacey D, Wulff K, Chikhalla N, Bernardo T. From FAIR to FAIRS: Data security by design for the global burden of animal diseases. Agron J. 2022 Sep;114(5):2693–9.

18. U.S. Department of Health & Human Services, Rights (OCR) O for C. The HIPAA Privacy Rule [Internet]. 2008 [cited 2024 Sep 17]. Available from: https://www.hhs.gov/hipaa/for-professionals/privacy/index.html

19. Afolayan AO, Bernal JF, Gayeta JM, Masim ML, Shamanna V, Abrudan M, et al. Overcoming Data Bottlenecks in Genomic Pathogen Surveillance. Clin Infect Dis. 2021 Dec 1;73(Supplement_4):S267–74.

20. Andrews S. FastQC: a quality control tool for high throughput sequence data. Cambridge, United Kingdom; 2010.

21. Chen S, Zhou Y, Chen Y, Gu J. fastp: an ultra-fast all-in-one FASTQ preprocessor. Bioinformatics. 2018 Sep 1;34(17):i884–90.

22. Seemann T. Shovill [Internet]. 2024 [cited 2024 Sep 17]. Available from: https://github.com/tseemann/shovill

23. Craddock HA, Motro Y, Zilberman B, Khalfin B, Bardenstein S, Moran-Gilad J. Long-Read Sequencing and Hybrid Assembly for Genomic Analysis of Clinical Brucella melitensis Isolates. Microorganisms. 2022 Mar 14;10(3):619.

24. Seemann T. any2fasta [Internet]. 2024 [cited 2024 Sep 17]. Available from: https://github.com/tseemann/any2fasta

25. Gurevich A, Saveliev V, Vyahhi N, Tesler G. QUAST: quality assessment tool for genome assemblies. Bioinformatics. 2013 Apr 15;29(8):1072–5.

26. Underwood A. BactInspector [Internet]. 2020 [cited 2024 Sep 17]. Available from: https://gitlab.com/antunderwood/bactinspector

27. Schoch CL, Ciufo S, Domrachev M, Hotton CL, Kannan S, Khovanskaya R, et al. NCBI Taxonomy: a comprehensive update on curation, resources and tools. Database [Internet]. 2020 Jan 1 [cited 2024 Sep 17];2020. Available from: https://academic.oup.com/database/article/doi/10.1093/database/baaa062/5881509

28. Seemann T. mlst [Internet]. 2024 [cited 2024 Sep 17]. Available from: https://github.com/tseemann/mlst

29. Feldgarden M, Brover V, Gonzalez-Escalona N, Frye JG, Haendiges J, Haft DH, et al. AMRFinderPlus and the Reference Gene Catalog facilitate examination of the genomic links among antimicrobial resistance, stress response, and virulence. Sci Rep. 2021 Jun 16;11(1).

30. Robertson J, Nash JHE. MOB-suite: software tools for clustering, reconstruction and typing of plasmids from draft assemblies. Microb Genomics. 2018 Aug 1;4(8).

31. Jain A, Singhal N, Kumar M. AFRbase: a database of protein mutations responsible for antifungal resistance. Martelli PL, editor. Bioinformatics. 2023 Nov 1;39(11).

32. Seemann T. Snippy [Internet]. 2024 [cited 2024 Sep 17]. Available from: https://github.com/tseemann/snippy

33. Harris SR. SKA: Split Kmer Analysis Toolkit for Bacterial Genomic Epidemiology [Internet]. Cold Spring Harbor Laboratory; 2018 [cited 2024 Sep 17]. Available from: http://biorxiv.org/lookup/doi/10.1101/453142

34. Page AJ, Taylor B, Delaney AJ, Soares J, Seemann T, Keane A, et al. SNP-sites: rapid efficient extraction of SNPs from multi-FASTA alignments. Microb Genomics.

35. Seemann T. snp-dists [Internet]. 2024 [cited 2024 Sep 17]. Available from: https://github.com/tseemann/snp-dists

36. Minh BQ, Schmidt HA, Chernomor O, Schrempf D, Woodhams MD, Von Haeseler A, et al. IQ-TREE 2: New Models and Efficient Methods for Phylogenetic Inference in the Genomic Era. Teeling E, editor. Mol Biol Evol. 2020 May 1;37(5):1530–4.

37. Sagulenko P, Puller V, Neher RA. TreeTime: Maximum-likelihood phylodynamic analysis. Virus Evol. 2018 Jan 1;4(1).

38. Hadfield J, Megill C, Bell SM, Huddleston J, Potter B, Callender C, et al. Nextstrain: real-time tracking of pathogen evolution. Kelso J, editor. Bioinformatics. 2018 Dec 1;34(23):4121–3.

39. Smith MR. Robust Analysis of Phylogenetic Tree Space. Hoehna S, editor. Syst Biol. 2022 Aug 10;71(5):1255–70.

40. Manara S, Pasolli E, Dolce D, Ravenni N, Campana S, Armanini F, et al. Whole-genome epidemiology, characterisation, and phylogenetic reconstruction of Staphylococcus aureus strains in a paediatric hospital. Genome Med [Internet]. 2018 Dec [cited 2024 Sep 17];10(1). Available from: https://genomemedicine.biomedcentral.com/articles/10.1186/s13073-018-0593-7

41. Eisenberger D, Tuschak C, Werner M, Bogdan C, Bollinger T, Hossain H, et al. Whole-genome analysis of vancomycin-resistant Enterococcus faecium causing nosocomial outbreaks suggests the occurrence of few endemic clonal lineages in Bavaria, Germany. J Antimicrob Chemother. 2020 Jun 1;75(6):1398–404.

42. Timme RE, Rand H, Shumway M, Trees EK, Simmons M, Agarwala R, et al. Benchmark datasets for phylogenomic pipeline validation, applications for foodborne pathogen surveillance. PeerJ. 2017 Oct 6;5:e3893.

43. Lockhart SR, Etienne KA, Vallabhaneni S, Farooqi J, Chowdhary A, Govender NP, et al. Simultaneous Emergence of Multidrug-Resistant *Candida auris* on 3 Continents Confirmed by Whole-Genome Sequencing and Epidemiological Analyses. Clin Infect Dis. 2017 Jan 15;64(2):134–40.

45. Huson DH, Scornavacca C. Dendroscope 3: An Interactive Tool for Rooted Phylogenetic Trees and Networks. Syst Biol. 2012 Dec 1;61(6):1061–7.

46. Smith MR. Information theoretic generalized Robinson–Foulds metrics for comparing phylogenetic trees. 2020;36(20):5007–13.

47. Harris PNA, Ben Zakour NL, Roberts LW, Wailan AM, Zowawi HM, Tambyah PA, et al. Whole genome analysis of cephalosporin-resistant Escherichia coli from bloodstream infections in Australia, New Zealand and Singapore: high prevalence of CMY-2 producers and ST131 carrying blaCTX-M-15 and blaCTX-M-27. J Antimicrob Chemother. 2018 Mar 1;73(3):634–42.

