## Supplementary Material 1 for "Solu – a cloud platform for real-time genomic pathogen surveillance"

### S1. Bioinformatics pipeline

### S1.A. Description of analyses ran for each sample

All tools are run using their default parameters unless otherwise specified.

**Supported input formats**

The pipeline supports three input formats: short reads as FASTQ, long reads as FASTQ, and assembled genomes as FASTA. Analysis of the long reads format is still considered experimental, and the methodology is subject to change. Compressed gzip archives (.gz) are also supported for all filetypes.

***De novo* assembly and standardization**

Short reads uploads are quality assured with FastQC v0.12. (1), pre-processed using fastp v0.23.4. (2) and assembled with Shovill v1.1.0 (3). Assembly files from all three sources – short reads, long reads, and direct user upload – are standardized using any2fasta v0.4.2. (4) to ensure consistency for downstream analyses. The parameters “-n” and “-u” of any2fasta are used in order to convert all unambiguous bases to “N” and all bases to uppercase. Unless otherwise noted, all downstream analyses are run uniformly for all upload formats, using the standardized assembly as their input. Assembly quality is assessed with QUAST v5.2.0 (5).

All downstream analyses use the standardized assembly as their input unless otherwise specified.

**Species identification**

Species are identified with Bactinspector v0.1.3 (6) against a database augmented with fungi. The augmented database contains the default BactInspector database, all fungal reference sequences from NCBI taxonomy (7), and clade-level references for *Candida auris*. The accessions used as references for *Candida auris* clades are: GCA_002759435 (clade I), GCA_003013715 (clade II), GCA_002775015 (clade III), GCA_003014415 (clade IV), GCA_016809505 (clade V), GCA_032367535 (clade VI).

**Genome size check**

After detecting the species, the assembled genome’s size is compared to the species’ expected genome size. The species-specific expected genome sizes are obtained from the NCBI genome size check API (8). Genome size check is implemented for a limited selection of species: *Acinetobacter baumannii, Acinetobacter variabilis, Acinetobacter pittii, Bacillus anthracis, Burkholderia cepacia, Burkholderia pseudomallei, Campylobacter coli, Campylobacter jejuni, Candida auris, Citrobacter freundii, Citrobacter portucalensis, Citrobacter werkmanii, Enterobacter asburiae, Enterobacter cloacae, Enterobacter hormaechei, Clostridium difficile, Clostridium perfringens, Enterococcus durans, Enterococcus faecalis, Enterococcus faecium, Enterococcus hirae, Escherichia coli, Haemophilus influenzae, Helicobacter pylori, Klebsiella aerogenes, Klebsiella oxytoca, Klebsiella pneumoniae, Klebsiella quasipneumoniae, Klebsiella variicola, Listeria monocytogenes, Mycobacteroides abscessus, Mycobacterium tuberculosis, Neisseria gonorrhoeae, Neisseria meningitidis, Proteus mirabilis, Pseudomonas aeruginosa, Salmonella enterica, Serratia marcescens, Shigella sonnei, Staphylococcus aureus, Staphylococcus epidermidis, Staphylococcus hominis, Staphylococcus pseudintermedius, Staphylococcus warneri, Staphylococcus xylosus, Stenotrophomonas maltophilia, Streptococcus agalactiae, Streptococcus mitis, Streptococcus pneumoniae, Streptococcus pyogenes, Vibrio cholerae, Vibrio parahaemolyticus, Vibrio vulfinicus*

**MLST (Multi-Locus Sequence Type)**

Genomes are queried against the PubMLST database to determine the Multi-Locus Sequence Type using mlst v2.23.0 (9). MLST is only run for bacterial species.

**Annotation of antimicrobial resistance (AMR) genes**

AMR and virulence genes of bacterial samples are annotated with AMRFinderPlus v3.11 (10), using “--organism” parameter for species-specific point mutations. In addition, the “--plus” parameter is used to detect genes impacting virulence or resistance to heat, biocide, metal, or acid.

In addition, annotation of antifungal resistance (AFR) genes is available for the species *Candida auris*. AFR loci are detected using the AMRFinderPlus tool with a custom database based on AFRBase (11). The current AFR database contains the following point mutations: *ERG11_E343D, ERG11_F126L, ERG11_F126T, ERG11_F444L, ERG11_I466M, ERG11_K143R, ERG11_K177R, ERG11_L376I, ERG11_N335S, ERG11_V125A, ERG11_Y132F, ERG11_Y501H, FKS1_S639F, FKS1_S639P, FUR1_F211I, CDR1_V704L.*

**Plasmid analysis**

Plasmid reconstruction and typing is performed using MOB-suite v3.1.9 (12). Plasmid analysis is only run for bacterial species.

### S1.B. Description of analyses between samples of the same species

The pipeline runs a phylogenetic comparison between all the uploaded samples that are detected as the same species. Phylogenetic comparison consists of pairwise SNP distances, clustering and phylogenetic tree inference, and is based on creating a multiple sequence alignment.

**Multiple sequence alignment**

Depending on the species, the pipeline has two different methods for creating a multiple sequence alignment. The primary method is based on aligning each sample to the species’ reference genome, and it is implemented for a set of commonly uploaded species. For the species to which the reference-based method is not yet available, a reference-free approach is used instead. The reference-based method is currently available for the following species: *Acinetobacter baumannii, Acinetobacter pittii, Candida auris, Cutibacterium acnes, Campylobacter coli, Enterobacter cloacae, Enterobacter hormaechei, Enterococcus faecium, Escherichia coli, Klebsiella quasipneumoniae, Klebsiella pneumoniae, Listeria monocytogenes, Mycobacteroides abscessus, Pseudomonas aeruginosa, Salmonella enterica, Serratia marcescens, Staphylococcus aureus, Staphylococcus epidermidis, Streptococcus pneumoniae, Vibrio cholerae.*

Reference-based alignment begins with aligning each sample to the species’ reference genome using Snippy v4.6.0 (13), with input parameters depending on the upload format. Samples uploaded as short paired-end reads are aligned using the “-R1” and “-R2” parameters, while samples uploaded as long reads or assembled genomes are aligned using Snippy’s “--contig” input option, which shreds the genome into pseudo-reads before alignment. The alignments of each sample are merged into a multiple sequence alignment using the “snippy core” command from Snippy. Low-quality SNPs are filtered from the multiple sequence alignment using an in-house Python script with the following logic. First, SNPs within 10 positions of another SNP are ignored to prune out recombination-related mutations. Second, SNPs within 15 positions of an ambiguous base (N) are ignored because they may contain sequencing or alignment errors.

Reference-free multiple sequence alignment is computed using the “ska align” command of SKA v1.0. (14) No separate SNP filtering is performed after the reference-free alignment.

After acquiring the multiple sequence alignment, the downstream phylogenetic comparison analyses are computed identically regardless of the alignment approach used.

**Phylogenetic comparison**

The pairwise SNP distances between each sample are computed from the multiple sequence alignment using snp-sites v.2.5.1 (15) and snp-dists v0.8.2 (16).

Samples are clustered using a 20-SNP single-linkage clustering threshold with an in-house Python script that utilizes the agglomerative clustering implementation from Scikit-learn library (17). The cluster names are automatically generated.

A phylogenetic tree for each species is inferred with the IQ-TREE v2.0.7 (18) maximum-likelihood algorithm. Prior to building the tree, constant sites are removed from the multiple sequence alignment using snp-sites v.2.5.1 (15). To preserve branch lengths, the number of removed sites per base is tracked using “snp-sites -C” and provided to IQ-TREE using the “-fconst” parameter. In addition, the parameter “-czb” parameter is used to collapse near-zero branches. The raw output from IQ-TREE is midpoint-rooted using TreeTime v0.11.2 (19). Both IQ-TREE and TreeTime are run using the Augur v24.2.2 toolkit (20).

### References

1.	Andrews S. FastQC: a quality control tool for high throughput sequence data. Cambridge, United Kingdom; 2010.

2.	Chen S, Zhou Y, Chen Y, Gu J. fastp: an ultra-fast all-in-one FASTQ preprocessor. Bioinformatics. 2018 Sep 1;34(17):i884–90.

3.	Seemann T. Shovill [Internet]. 2024 [cited 2024 Sep 17]. Available from: https://github.com/tseemann/shovill

4.	Seemann T. any2fasta [Internet]. 2024 [cited 2024 Sep 17]. Available from: https://github.com/tseemann/any2fasta

5.	Gurevich A, Saveliev V, Vyahhi N, Tesler G. QUAST: quality assessment tool for genome assemblies. Bioinformatics. 2013 Apr 15;29(8):1072–5.

6.	Underwood A. BactInspector [Internet]. 2020 [cited 2024 Sep 17]. Available from: https://gitlab.com/antunderwood/bactinspector

7.	Schoch CL, Ciufo S, Domrachev M, Hotton CL, Kannan S, Khovanskaya R, et al. NCBI Taxonomy: a comprehensive update on curation, resources and tools. Database [Internet]. 2020 Jan 1 [cited 2024 Sep 17];2020. Available from: https://academic.oup.com/database/article/doi/10.1093/database/baaa062/5881509

8.	NCBI Genome Size Check [Internet]. [cited 2024 Sep 17]. Available from: https://www.ncbi.nlm.nih.gov/genbank/genome-size-check/

9.	Seemann T. mlst [Internet]. 2024 [cited 2024 Sep 17]. Available from: https://github.com/tseemann/mlst

10.	Feldgarden M, Brover V, Gonzalez-Escalona N, Frye JG, Haendiges J, Haft DH, et al. AMRFinderPlus and the Reference Gene Catalog facilitate examination of the genomic links among antimicrobial resistance, stress response, and virulence. Sci Rep. 2021 Jun 16;11(1).

11.	Jain A, Singhal N, Kumar M. AFRbase: a database of protein mutations responsible for antifungal resistance. Martelli PL, editor. Bioinformatics. 2023 Nov 1;39(11).

12.	Robertson J, Nash JHE. MOB-suite: software tools for clustering, reconstruction and typing of plasmids from draft assemblies. Microb Genomics. 2018 Aug 1;4(8).

13.	Seemann T. Snippy [Internet]. 2024 [cited 2024 Sep 17]. Available from: https://github.com/tseemann/snippy

14.	Harris SR. SKA: Split Kmer Analysis Toolkit for Bacterial Genomic Epidemiology [Internet]. Cold Spring Harbor Laboratory; 2018 [cited 2024 Sep 17]. Available from: http://biorxiv.org/lookup/doi/10.1101/453142

15.	Page AJ, Taylor B, Delaney AJ, Soares J, Seemann T, Keane A, et al. SNP-sites: rapid efficient extraction of SNPs from multi- FASTA alignments. Microb Genomics.

16.	Seemann T. snp-dists [Internet]. 2024 [cited 2024 Sep 17]. Available from: https://github.com/tseemann/snp-dists

17.	Pedregosa F, Varoquaux G, Gramfort A, Michel V, Thirion B, Grisel O, et al. Scikit-learn: Machine Learning in Python. Mach Learn PYTHON. 2011;12(85):2825–30.

18.	Minh BQ, Schmidt HA, Chernomor O, Schrempf D, Woodhams MD, Von Haeseler A, et al. IQ-TREE 2: New Models and Efficient Methods for Phylogenetic Inference in the Genomic Era. Teeling E, editor. Mol Biol Evol. 2020 May 1;37(5):1530–4.

19.	Sagulenko P, Puller V, Neher RA. TreeTime: Maximum-likelihood phylodynamic analysis. Virus Evol. 2018 Jan 1;4(1).

20.	Hadfield J, Megill C, Bell SM, Huddleston J, Potter B, Callender C, et al. Nextstrain: real-time tracking of pathogen evolution. Kelso J, editor. Bioinformatics. 2018 Dec 1;34(23):4121–3.
