## Supplementary Material 2 for "Solu – a cloud platform for real-time genomic pathogen surveillance"

### S2. Additional phylogenetics results

**Figure 1.** Solu’s phylogram of the S. aureus dataset vs. reported sequence types (outer ring)

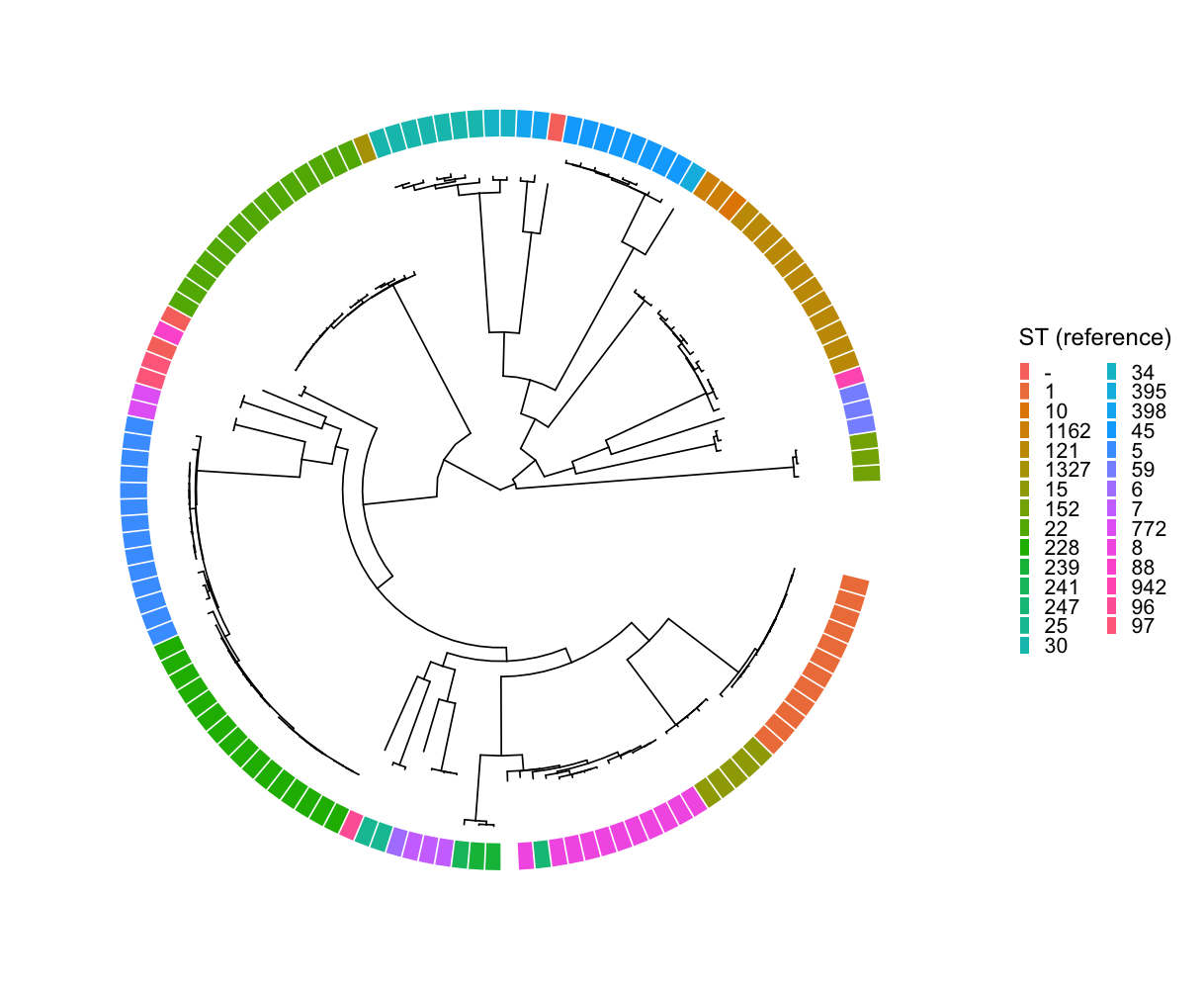

**Figure 2**. Solu’s phylogram of the Candida auris dataset vs. reported clade (outer ring)

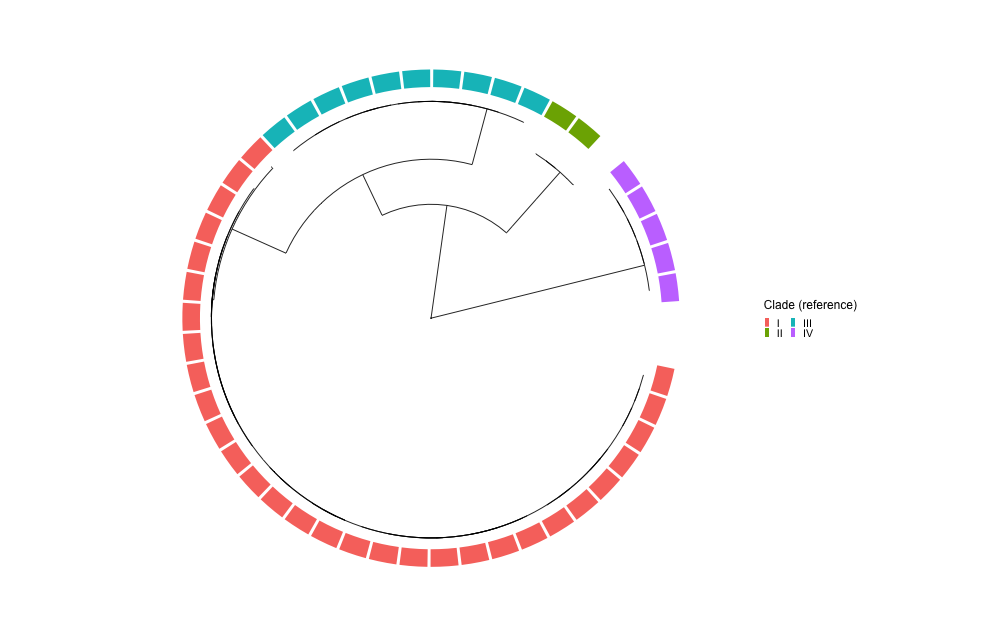

**Table 1.** Validation of Solu’s reference-based vs. reference-free phylogenetics

| **Dataset** | **Robinson-Foulds distance of Solu’s reference-based tree vs. Solu’s reference-free tree** |
| --- | --- |
| *S. aureus* | 0.08093671 |
| *E. faecium* | 0.2694115 |
| *S. enterica* | 0.4644507 |
| *C. auris* | 0.2362036 |

The tree distances were calculated with TreeDist 2.9.1 (1).

**Figure 3**. Tanglegram of *Staphylococcus aureus* reference-free (left) vs. reference-based (right) phylogenetic trees
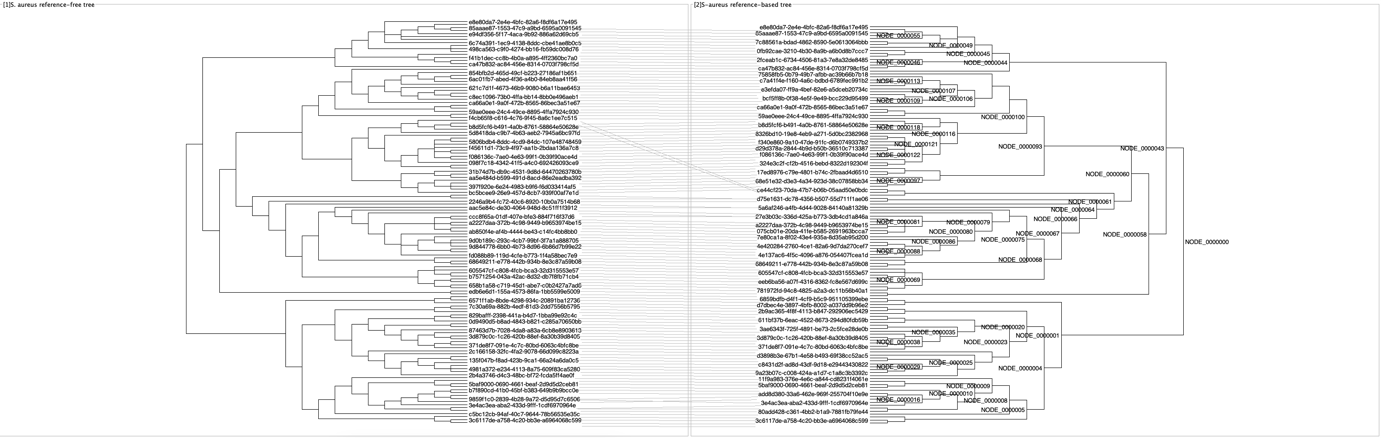

**Figure 4**. Tanglegram of *Salmonella enterica* reference-free (left) vs. reference-based (right) phylogenetic trees
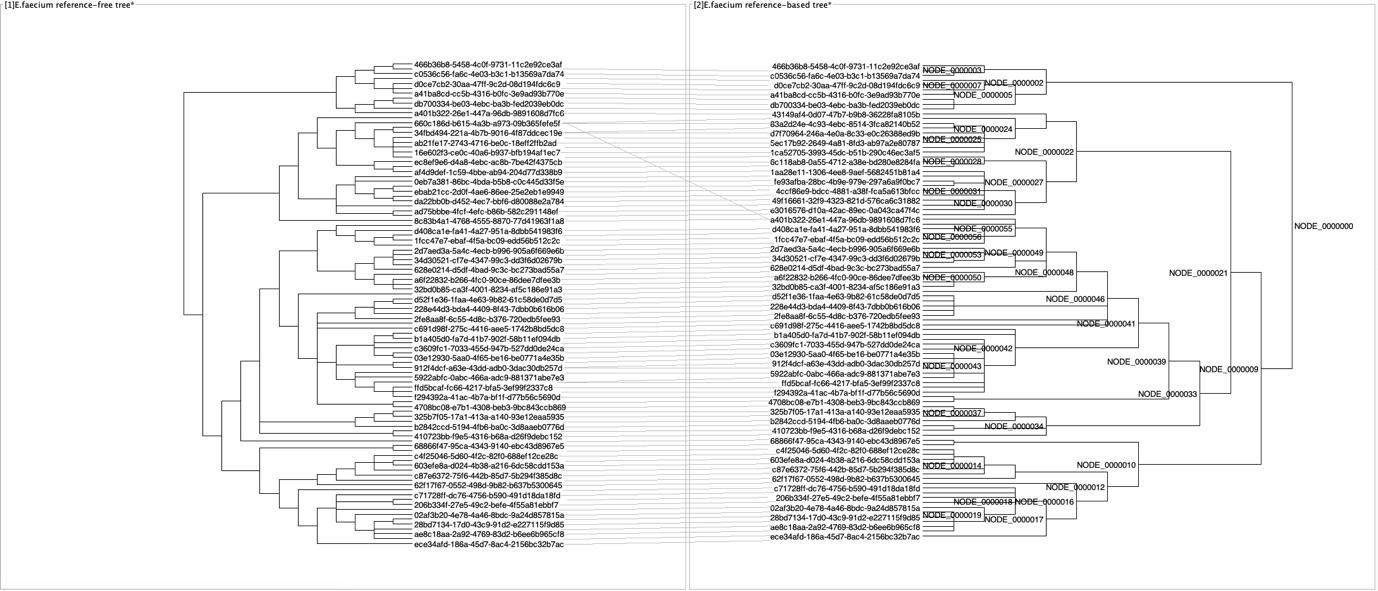

### **Figure 5**. Tanglegram of *Candida auris* reference-free (left) vs. reference-based (right) phylogenetic trees

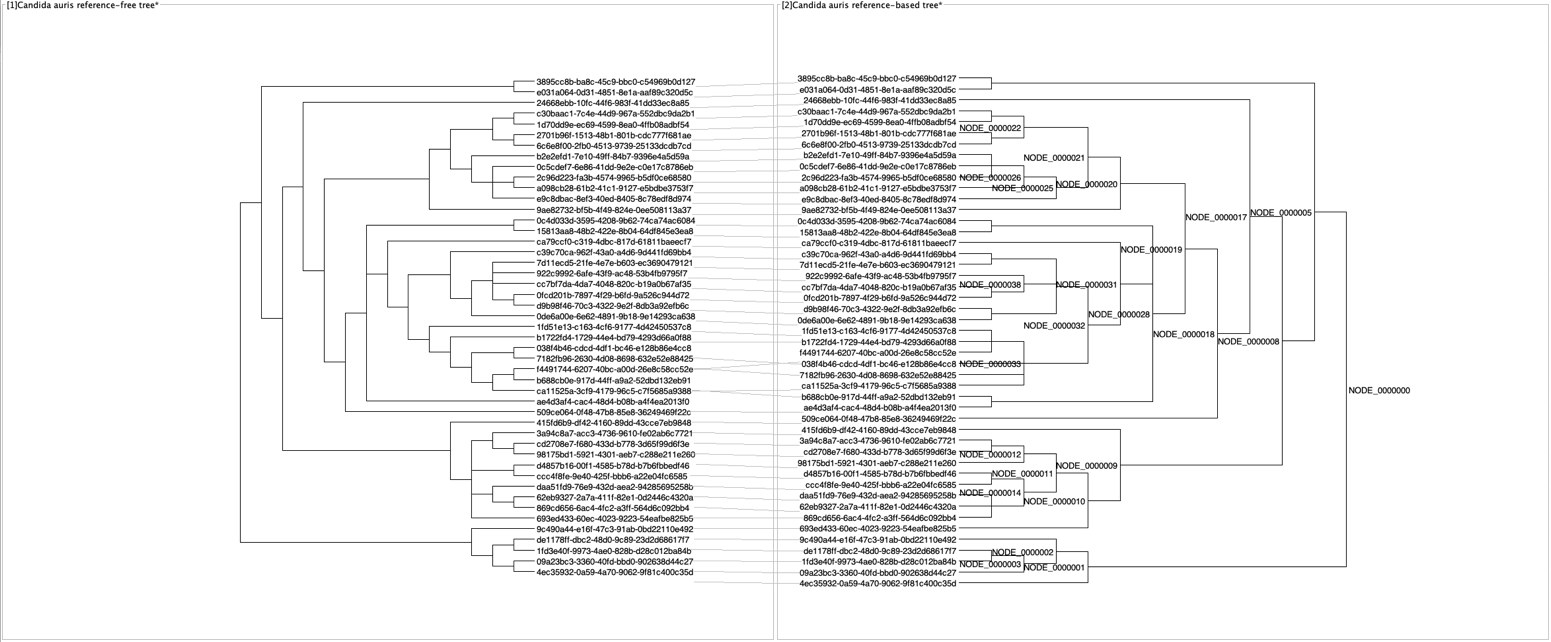

### **Figure 6**. Tanglegram of *Salmonella enterica* reference-free (left) vs. reference-based (right) phylogenetic trees

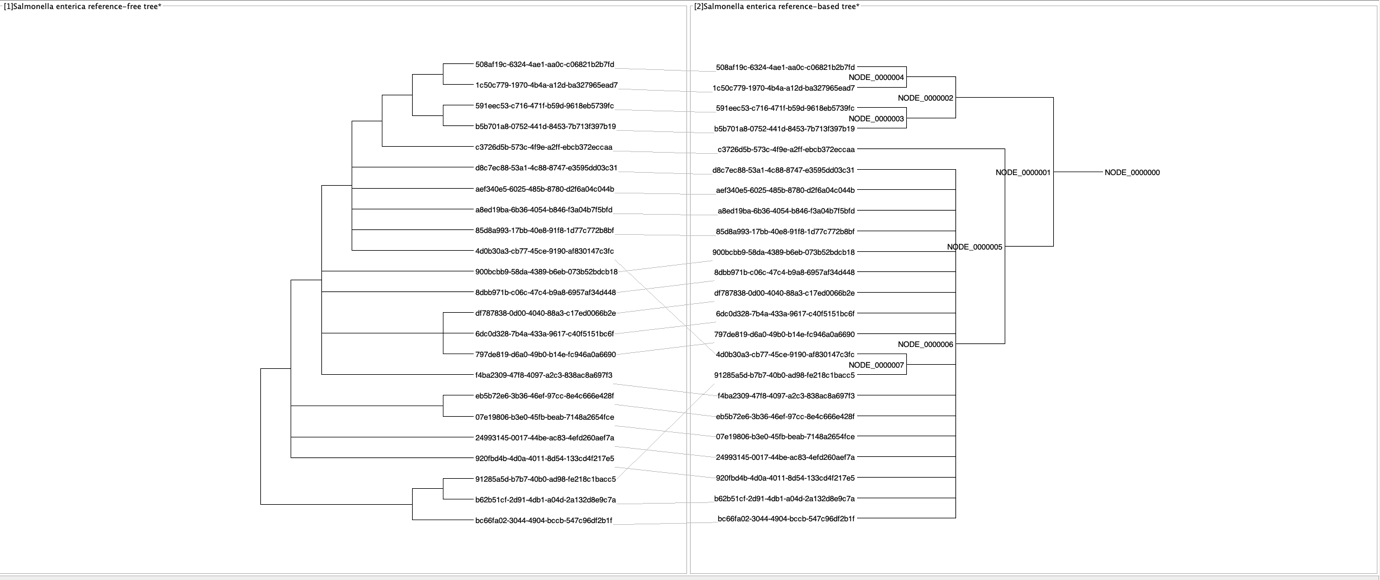

All tanglegrams were created with Dendroscope v 3.8.10 (2).

### References

1.	Smith MR. Information theoretic generalized Robinson–Foulds metrics for comparing phylogenetic trees. 2020;36(20):5007–13.

2.	Huson DH, Scornavacca C. Dendroscope 3: An Interactive Tool for Rooted Phylogenetic Trees and Networks. Syst Biol. 2012 Dec 1;61(6):1061–7.
